# AlphaDesign: A *de novo* protein design framework based on AlphaFold

**DOI:** 10.1101/2021.10.11.463937

**Authors:** Michael Jendrusch, Jan O. Korbel, S. Kashif Sadiq

**Affiliations:** Genome Biology Unit, European Molecular Biology Laboratory (EMBL), Meyerhofstrasse 1, Heidelberg 69117, Germany

**Keywords:** AlphaFold, computational protein design, self-assembly, biologics, network inversion, evolutionary algorithms, computational biophysics

## Abstract

*De novo* protein design is a longstanding fundamental goal of synthetic biology, but has been hindered by the difficulty in reliable prediction of accurate high-resolution protein structures from sequence. Recent advances in the accuracy of protein structure prediction methods, such as AlphaFold (AF), have facilitated proteome scale structural predictions of monomeric proteins. Here we develop AlphaDesign, a computational framework for *de novo* protein design that embeds AF as an oracle within an optimisable design process. Our framework enables rapid prediction of completely novel protein monomers starting from random sequences. These are shown to adopt a diverse array of folds within the known protein space. A recent and unexpected utility of AF to predict the structure of protein complexes, further allows our framework to design higher-order complexes. Subsequently a range of predictions are made for monomers, homodimers, heterodimers as well as higher-order homo-oligomers - trimers to hexamers. Our analyses also show potential for designing proteins that bind to a pre-specified target protein. Structural integrity of predicted structures is validated and confirmed by standard *ab initio* folding and structural analysis methods as well as more extensively by performing rigorous all-atom molecular dynamics simulations and analysing the corresponding structural flexibility, intramonomer and interfacial amino-acid contacts. These analyses demonstrate widespread maintenance of structural integrity and suggests that our framework allows for fairly accurate protein design. Strikingly, our approach also reveals the capacity of AF to predict proteins that switch conformation upon complex formation, such as involving switches from *α*-helices to *β*-sheets during amyloid filament formation. Correspondingly, when integrated into our design framework, our approach reveals *de novo* design of a subset of proteins that switch conformation between monomeric and oligomeric state.

## I. INTRODUCTION

Proteins are the workhorses of the vast majority of life processes at the cellular scale. They carry out a myriad of functions ranging from the catalysis of biochemical reactions, mechanical functions involved in cell motility and the formation of sub-cellular architecture amongst many others. A central paradigm to understand their function has been based on the observation that proteins fold into complex, yet specific three-dimensional structures that vary depending on their amino-acid sequence, hypothesized to be their lowest energy state [1]. Thus experimental structure determination has been a primary pursuit of biology for the last 50 years [2] and given rise to over 10^5^ distinctly solved structures [3]. These have revealed a diverse array of fold topologies and geometries of individual proteins, classified into 41 architectures with 1390 fold topologies [4] and that 51% of structures previously extracted [5] from the protein data bank (PDB) [3] form quaternary structure through oligomer and complex formation [5].

A central driving question has been to what extent amino-acid sequence encodes structure and whether it is then possible to predict structure from sequence. This has fuelled a plethora of diverse computational protein structure prediction approaches across decades of effort [6–14]. The quest for improved methods has been embodied in an independently assessed biennial community-wide competition (CASP) that ranks method accuracy of participants’ approaches with respect to determined but unreleased structures resulting in steady progress towards predictive accuracy [15]. Recently, this has culminated in the development of AlphaFold [16] - hereafter, termed AF, a state-of-the-art neural network-based approach that has achieved comparable atomic accuracy with respect to crystal structure. AF has been applied at a proteome-wide scale across several species, resulting in a new database of predicted structures [17]. More recently, a community-wide assessment has revealed several important applications of AF ranging from amino acid variant effect prediction to cryo-EM model building [18]. Whilst intended for single-chain structure prediction, an unexpected consequence of AF’s input protocol allows prediction of non-contiguous chains, thus enabling prediction of protein complexes using the existing trained network [18, 19]. This feature has also been applied at proteome-wide scale to reveal novel core eukaryotic complexes [20]. These efforts culminated in the release of AlphaFold-Multimer [21], a version of AF explicitly trained for complex prediction.

Despite these advances, the protein folding problem remains far more complex than structure prediction alone. Proteins are not rigid structures, especially at physiological conditions. Firstly, proteins exhibit thermodynamic equilibrium between folded and unfolded forms [1]. Secondly, many proteins also undergo conformational changes with a well-described equilibrium between states, making use of these changes to enact function [22]. Moreover, there is an abundance of intrinsically disordered proteins (IDPs) - those that do not exhibit stable tertiary structures, or transiently fold depending on environmental context [22–25]. One notable biomedical example being the misfolding of amyloid-*α* helices into *β*-sheeted filaments associated with Alzheimer’s disease [26]. Intriguingly, metamorphic proteins - those that form multiple, stable, yet different folds - have also been discovered [27].

From a biophysical perspective, protein folding is made comprehensible using the concept of a folding energy funnel within the atomic configuration space [28]. Multiple energy minima then correspond to alternate stable states that make transitions with characterisable rates. Based on this concept, computational physics methods such as molecular dynamics (MD) simulations have provided a route to characterise the dynamics, thermodynamics and kinetics of conformational transitions [29, 30], *ab initio* folding [31], disordered transitions [32, 33] as well as protein-protein [34, 35] and protein-ligand [36] binding.

Notwithstanding the already-mentioned complex structure and dynamics of naturally occurring proteins, evolution has still only explored an infinitesimal portion of the potential protein sequence landscape [37]. There is therefore enormous potential in unlocking the fundamental biophysical principles of protein folding to design and engineer novel proteins that can exploit this vast space. Recent examples include protein logic gates [38], self-assembling systems [39] and targeted therapeutics [40]. Established methods for protein engineering have until recently focused on tuning naturally occurring proteins through iterative experimental selection processes such as directed evolution [41–43]. More recently, computational design approaches have enabled *de novo* protein design, encompassing a full suite of functionalities ranging from rules for topology selection, protein backbone construction, sequence optimisation as well combinations of these approaches [37, 44].

The current *de facto* standard in computational protein design is embodied by the Rosetta suite of protein design tools [45, 46] and Rosetta Remodel [47] in particular. It combines tools for all steps from selecting a desired protein topology to designing and validating a folding protein sequence. Protein design within Rosetta Remodel is composed of a combination of four tasks: topology specification, backbone generation and fixed-backbone design [47]. After specifying a desired protein topology a backbone structure can be generated using matching fragments extracted from existing proteins [45, 47]. Fragments matching a desired secondary structure and set of contacts are selected from PDB structures. Selected fragments are then sampled using Markov-Chain Monte Carlo to arrive at a pool of candidate backbone geometries [47]. Candidate backbone structures can then be equipped with a sequence using fixed-backbone protein design, optionally optimising the backbone between design steps [48]. Given a desired backbone geometry, the goal is to find an amino acid sequence which will fold that structure. Here, the Rosetta Design protocol [48] starts by populating the desired backbone with an all-valine sequence and runs Markov-Chain Monte Carlo to arrive at a low-energy sequence. The general strategy of Rosetta-based protein design has been successfully applied to a variety of design problems. These applications range from the first *de novo* designed proteins [48] to synthetic vaccines [49, 50], complex assemblies [51, 52] and enzymes [53, 54]. However, Markov-Chain Monte Carlo in structure space can be time-consuming and computationally demanding.

To tackle this shortcoming of classical protein design, there has been an increase in approaches applying neural networks to various design problems. Several works train generative models to directly generate protein sequences with a desired function [55–57]. [55] have trained a language model on the UniProt sequence database [58] to generate sequences with a specified function. [59] have expanded on [55] by additionally fine-tuning the generative model on a specific protein family or function. Beyond language models, other types of generative models, such as variational autoencoders [56] and generative adversarial networks [57] have been explored for direct protein sequence generation. While these approaches have been successful in generating protein sequences associated with a given function, they do not explicitly take into account structural information and thus cannot be applied to protein design tasks involving constraints on tertiary structure.

On the other side of the spectrum of neural network-based protein design, generative models have been explored for protein structure generation [60–62]. [60, 63] have trained generative adversarial networks to generate realistic backbone distance maps and coordinates. [62] have achieved the same goal using variational autoencoders. Both approaches have demonstrated fine-grained control over designed backbone structures by latent variable manipulation. While these approaches are a viable alternative to classical backbone design, they offer no guarantees on the designability of their predicted backbones. In contrast, our approach produces backbones and sequences which are by construction designable and high-confidence under AF.

Bridging the gap between structure-only and sequence-only approaches, [64–67] have trained neural networks on the PDB database of protein structures to predict protein sequence given a fixed backbone structure. These approaches rely on network components taking into account the geometry of the protein backbone together with the protein sequence. [64, 65, 67] have used per-residue local coordinate systems to reason about protein geometry, while [66] use network architecture adapted to Euclidean coordinate data. [68] have framed fixed-backbone design as a constraint satisfaction problem and have trained their network as a constraint solver. These methods require a neural network trained for a specific protein design task – fixed-backbone protein design – and by themselves cannot easily be extended to other design applications without retraining.

A number of works explore re-using previously trained predictors of protein structure or function as parts of an *in silico* screening framework [69–76]. In these approaches, a neural network is treated as a score function to evaluate the quality of protein designs. Designs are then improved using gradient-based [71–73], gradient-free [69, 74, 76] or neural-network based [75] optimisation.

[70] were the first to use an optimisation loop incorporating a neural network for structure prediction for *de novo* protein design. Their approach has since been extended to fixed-backbone design [71, 73] and protein scaffold generation for protein motif stabilisation [72]. [71, 72] use both Markov-Chain Monte Carlo and gradient descent as their means for optimisation. All of these approaches have made use of trRosetta [77] as a structure predictor. With the new release of AF, [74] have used it as a structure predictor for fixed-backbone protein design using greedy optimisation, with sequences initialised from a trained model. We expand previous approaches by constructing a flexible family of target functions for optimisation using AF [16] and extending the range of possible design tasks.

In this work, we develop a flexible framework for protein design by sequence optimisation using evolutionary algorithms. We embed AF [16] into a design loop as a structure prediction oracle. Our framework extends previous work on protein design using structure predictors [70, 71] by defining a flexible family of target functions encoding various protein design tasks and integrate this design loop with extensive validation using both Rosetta [45] and molecular dynamics simulations. We apply our framework to design de novo protein monomers, dimers and oligomers. We further design binders for target proteins using only their sequence, as well as design proteins which change conformation upon complex formation.

## II. *DE NOVO* DESIGN FRAMEWORK DEVELOPMENT

We frame protein design as a search problem to find the set of protein sequences for which a certain target function exceeds a fixed threshold. To take into account both sequence properties and all-atom protein structure, we integrate AF [16] into our target functions to provide high-quality structure prediction and measures of prediction confidence. We combine this with state-of-the art validation using Rosetta *ab initio* structure prediction [45] and molecular dynamics simulations. In the following we describe the components of our framework.

### A. Optimisation loop

Our approach to *de novo* protein design follows prior work using trRosetta [70, 71, 77]. We set up an optimisation loop which continuously mutates a pool of sequences, scores them using a neural network-based cost function and updates the sequence pool with the mutated sequences and scores (Fig. 1 A). In contrast to previous approaches using trRosetta, we use AF as a structure prediction oracle [16] and define our optimisation target as a flexible function of protein sequence, all-atom structure and AF confidence.

**FIG. 1.**
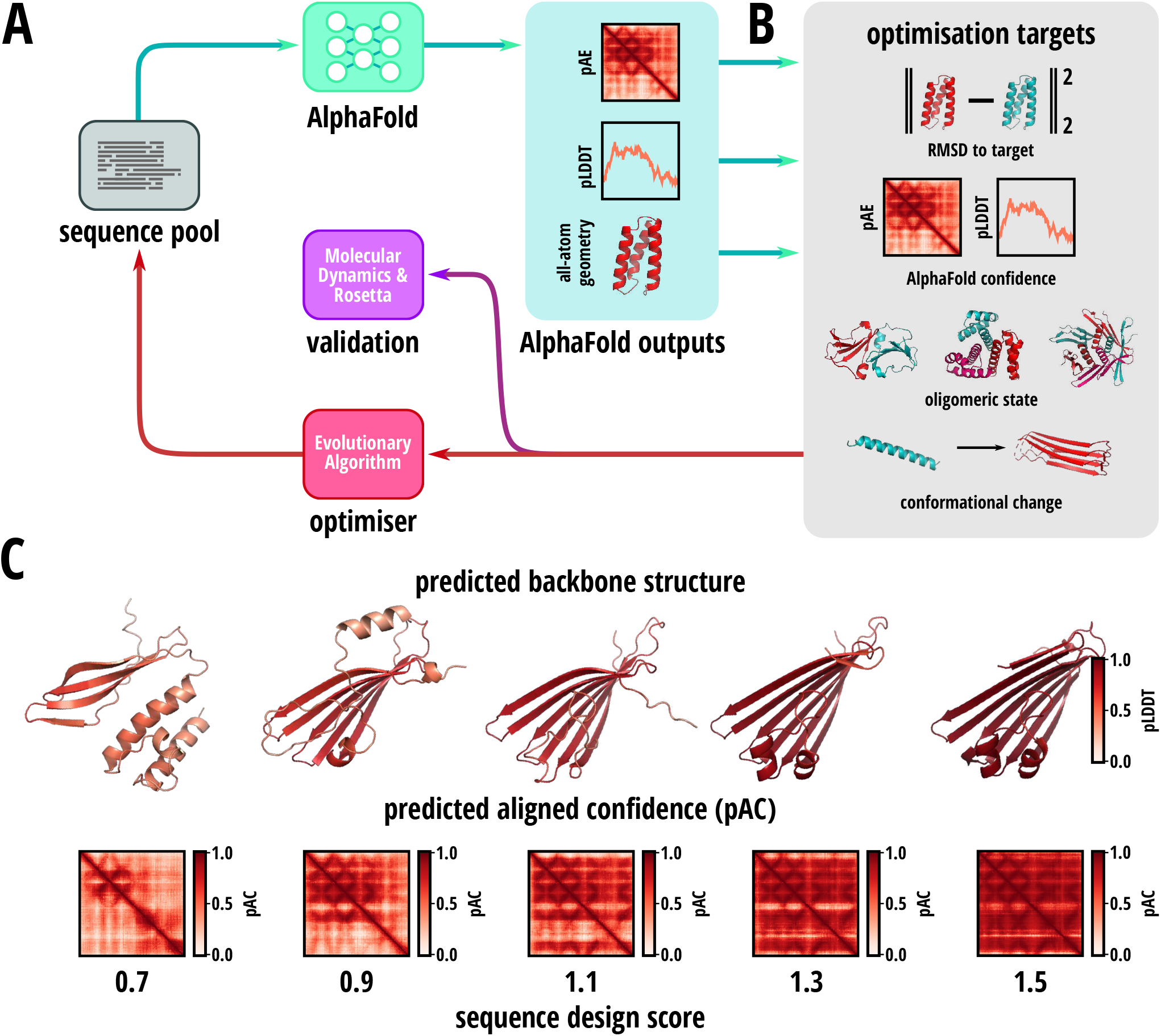
Method overview. **(A)** Core optimisation loop of AlphaDesign. A pool of amino acid sequences (grey) is maintained throughout the design process. AlphaFold (blue) predicts the all-atom structure and confidence metrics (pLDDT, pAC) for each sequence (light blue). These are combined into a target function maximised in the optimisation loop. The sequences and their target values are fed to an optimiser (red), which updates the sequence pool. If the target value exceeds a desired threshold, the sequence and its structure are submitted for validation using Rosetta and molecular dynamics simulations (purple). **(B)** Target function components. The optimisation target may be any function of the amino acid sequence, predicted all-atom structure and confidence metrics (pLDDT, pAC). Examples include RMSD to a target structure, maximisation of confidence, complex formation and conformational change upon complex formation. **(C)** Snapshots of a sequence at increasing target values during optimisation. For increasing target values from 0.7 to 1.5 the predicted structure (top) changes. Simultaneously, local confidence for each amino acid (pLDDT, top) and predicted aligned confidence for each pair of amino acids (pAC, bottom) increase.

AF is designed to process a sequence together with a corresponding multiple sequence alignment, it is possible to predict the structure for a single sequence by constructing an alignment containing that sequence alone [19]. While this decreases the capability of AF to predict correct structures with default parameters, previous work shows that increasing the number of AF iterations on a single input (referred to as recycling steps) can rescue prediction quality [19]. AF returns an all-atom structure for the input protein sequence together with predicted confidence measures (Fig. 1 A, AlphaFold outputs). AF confidence is expressed as a combination of the predicted local distance difference test (pLDDT) [78] and the predicted aligned error (pAE) [16]. pLDDT measures local model quality, while pAE provides a measure of confidence for each amino acid pair. We may then optimise sequences to maximise any target function 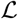 of these outputs. These can include minimising the RMSD to a target structure and maximising AF prediction confidence. As AF is suitable for complex structure prediction [18], target functions can also constrain oligomeric state or conformational change upon protein binding (Fig. 1 B). In general, any function of protein structure, sequence and prediction confidence may be used for optimisation.

For each sequence in the sequence pool we compute the value of the target function. Sequences are further recombined and mutated by an optimiser to explore sequence space. The sequence pool is updated with the mutated sequences. In this work, we use a simple evolutionary algorithm following [79]. However, other gradient-free or gradient-based optimisers can be substituted as desired. Throughout optimisation a protein changes its structure and both local and global confidence measures increase (Fig. 1 C). For a given protein design task, a threshold is selected above which a sequence is considered fully optimised. We return sequences above this threshold and submit a subset of them to validation using molecular dynamics simulations and Rosetta *ab initio* structure prediction [45].

### B. Target functions

When using gradient-free methods, we can use any function of the protein sequence, all-atom structure and AF confidence as the optimisation target. This opens up many possibilities for designing proteins and protein complexes with specific properties. In this work, we focus on cost functions making use of AF confidence and protein backbone geometry.

#### a. Confidence

AF models return two main measures of confidence for a protein structure prediction. These are the predicted local distance difference test (pLDDT) [78] and the predicted aligned error (pAE) [16]. pLDDT provides a measure of local confidence for each amino acid, while pAE provides the predicted error of each amino acid position in the local coordinate frame of each other amino acid. For both confidence measures, AF predicts a binned distribution of values. Instead of working directly with pAE values, we instead convert to predicted aligned confidence (pAC), which translates to *pAC* =1 – *μ_pAE_*/*pAE_max_* where *μ_pAE_* is the mean pAE and *pAE_max_* is the center of the highest pAE bin. Alternatively, we can work with the predicted template matching score (pTM) [16, 80].

All target functions in this work include a term maximising confidence in the form of:

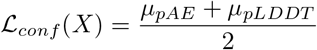

where *X* = (*x*, *pAE*, *pLDDT*) is a tuple containing the all-atom structure *x*, as well as the pAE and pLDDT for each protein.

#### b. Geometry

As AF returns protein all-atom coordinates [16], we may introduce target terms which depend on arbitrary geometrical features of a protein. The main building block for such terms are measures of difference between two sets of coordinates. We have implemented the following protein structure metrics to use within cost functions:

- Aligned error [16]:

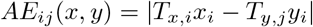

where *T_x,i_* is the coordinate transformation moving a vector to the local coordinate system at backbone atom *x_i_*. This is the aligned error used in AF training [16].
- Frame-Aligned Point Error [16]:

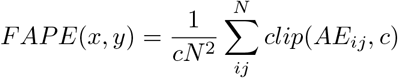

where *c* is a cutoff for the maximum error. This is the FAPE loss used in AF training.
- distance RMSD:

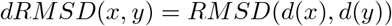
- Template Matching score [80]:

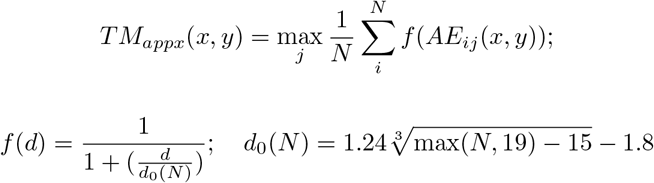

This is an approximation of the template matching score which does not require structural alignment [16]. This is our preferred measure for structural similarity, as it is normalised to 1 and provides a smoother target for highly dissimilar structures.

Furthermore, we can enforce general shape constraints for proteins using a compactness target 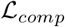, which penalises large protein radii:

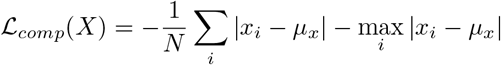

where *μ_x_* denotes the center of mass of the coordinates *x*.

#### c. Monomers

To generate globular monomeric proteins, we combine the confidence target with a weighted compactness target as follows:

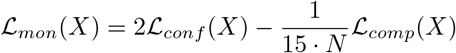

This trades off compactness for confidence in the case of elongated structures.

#### d. Complexes

For protein complex design, we use the same target function 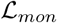 for each constituent monomer. This ensures each monomer has a stable high-confidence structure. Previous work identified inter-monomer pAE as a predictor of complex formation [19]. Low inter-monomer pAE corresponds to a high confidence in complex prediction. Therefore, we also impose the same confidence and compactness target on the complex, resulting in:

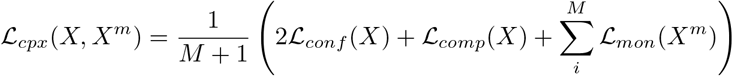

where *X* denote the AF predictions for the complex and *X^m^* denote the same for each monomer *m*.

#### e. Conformational change

To steer towards conformational change upon complex formation, we introduce an additional term to our oligomer cost function:

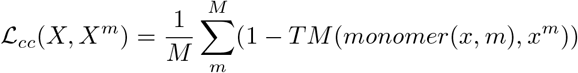

where *monomer*(*x*, *m*) extracts the coordinates of monomer m from the structure of the oligomer *x*. Essentially, maximising this target function minimises the TM score [80] between a protein structure as part of a complex and its structure as a monomer.

### C. Input representation

We represent amino acid sequences as one-hot encoded arrays with 20 classes, one for each standard amino acid. This allows us to use both gradient-free and gradient-based optimisers, as we can estimate the gradient through a one-hot representation using a straight-through estimator or similar [81]. As input to AF, we construct an additional multiple-sequence alignment with only the single input sequence, as well as blank template features.

#### a. Sequence templates

To reduce search space size and implement exact sequence constraints without introducing additional cost functions, we represent optimised sequences as a pair (*S*, *T*) where 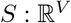 is a one-hot representation of all *V* variable amino-acids and 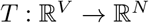 is a template mapping which assembles the *V* variable amino acids into the final evaluated protein sequence of length *N*. For example, to design a homodimer with monomer size 64, the actual number of variable amino acids is *V* = 64 and the template *T* simply concatenates the input sequence 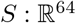 with itself.

#### b. Complex representation

Following [19], we represent chain breaks for complex prediction by introducing an increment greater than 32 to the residue indices of each additional chain beyond the first. We use this increment to separate all residues on different chains by more than 32, which is the maximum relative residue index difference separately embedded by AF [16]. In addition, following [19] we split the multiple-sequence alignment features, such that each monomer has a separate copy of its sequence alignment features, with gaps at the sequence positions of other monomers.

## III. MATERIALS AND METHODS

### A. Prediction studies

Complex and conformational change predictions were performed using Colabfold [19] using the advanced notebook. In all cases, all five AF parameter sets were used, and models were selected by highest predicted template matching score [80]. Complex queries were predicted without paired MSA. The resulting top model was structurally aligned using PyMol [82] for RMSD reporting. All prediction runs are summarised in Table S1.

### B. Design studies

#### a. Optimisation

For optimisation an evolution strategy optimiser following [79] was used. Population size was set as 10. During mutation population size was expanded by a factor of 2. Sequences with a suboptimality of at most 10% were considered for mutation and recombination. Recombination was applied by crossover with probability of 10% at each sequence position. Optimisation was considered complete for sequences with 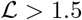 for 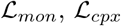 and 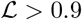 for target functions including 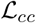. Tab. I and Tab. II contain further parameters of optimisation runs performed.

**TABLE I.** *De novo* protein design runs and parameters.

| name | sequence length | $N$ monomers | homooligomer | $N$ recycling | target function |
| --- | --- | --- | --- | --- | --- |
| monomers 32 | 32 | 1 | – | 4 | $\mathcal{L}_{mon}$ |
| monomers 64 | 64 | 1 | – | 4 | $\mathcal{L}_{mon}$ |
| monomers 128 | 128 | 1 | – | 4 | $\mathcal{L}_{mon}$ |
| monomers 128 | 128 | 1 | – | 4 | $\mathcal{L}_{mon}$ |
| homodimers 32 | 32 | 2 | yes | 4 | $\mathcal{L}_{cpx}$ |
| homodimers 64 | 64 | 2 | yes | 4 | $\mathcal{L}_{cpx}$ |
| homodimers 128 | 128 | 2 | yes | 2 | $\mathcal{L}_{cpx}$ |
| heterodimers 32 | 32 | 2 | no | 4 | $\mathcal{L}_{cpx}$ |
| heterodimers 64 | 64 | 2 | no | 4 | $\mathcal{L}_{cpx}$ |
| heterodimers 128 | 128 | 2 | no | 2 | $\mathcal{L}_{cpx}$ |
| trimers 32 nr4 | 32 | 3 | yes | 4 | $\mathcal{L}_{cpx}$ |
| trimers 64 nr4 | 64 | 3 | yes | 4 | $\mathcal{L}_{cpx}$ |
| trimers 32 nr2 | 32 | 3 | yes | 2 | $\mathcal{L}_{cpx}$ |
| trimers 64 nr2 | 64 | 3 | yes | 2 | $\mathcal{L}_{cpx}$ |
| tetramers 32 nr4 | 32 | 4 | yes | 4 | $\mathcal{L}_{cpx}$ |
| tetramers 64 nr4 | 64 | 4 | yes | 4 | $\mathcal{L}_{cpx}$ |
| tetramers 32 nr2 | 32 | 4 | yes | 2 | $\mathcal{L}_{cpx}$ |
| tetramers 64 nr2 | 64 | 4 | yes | 2 | $\mathcal{L}_{cpx}$ |
| pentamers 32 nr4 | 32 | 5 | yes | 4 | $\mathcal{L}_{cpx}$ |
| pentamers 64 nr4 | 64 | 5 | yes | 4 | $\mathcal{L}_{cpx}$ |
| pentamers 32 nr2 | 32 | 5 | yes | 2 | $\mathcal{L}_{cpx}$ |
| pentamers 64 nr2 | 64 | 5 | yes | 2 | $\mathcal{L}_{cpx}$ |
| conformation change dimers 32 | 32 | 2 | no | 4 | $0.7 \cdot \mathcal{L}_{cpx} + 0.3 \cdot \mathcal{L}_{cc}$ |
| conformation change dimers 64 | 64 | 2 | no | 4 | $0.7 \cdot \mathcal{L}_{cpx} + 0.3 \cdot \mathcal{L}_{cc}$ |
| conformation change trimers 32 | 32 | 3 | yes | 4 | $0.7 \cdot \mathcal{L}_{cpx} + 0.3 \cdot \mathcal{L}_{cc}$ |

**TABLE II.** Binder design design runs and parameters.

| name | binder length | target | $N$ recycling | target function |
| --- | --- | --- | --- | --- |
| binders 64-64-0 | 64 | monomers 64 result 0 | 4 | $\mathcal{L}_{cpx}$ |
| binders 64-64-2 | 128 | monomers 64 result 2 | 4 | $\mathcal{L}_{cpx}$ |

#### b. AlphaFold configuration

For optimisation, AF was configured for single-sequence use by disabling ensembling, templates, extra MSA features and restricting the number of MSA features to the number of monomers modelled. The number of AF iterations (recycling steps) was kept as a parameter for each optimisation run (Tab. I, Tab. II). For larger protein complexes, the number of iterations was decreased to 2 to speed up computation. The parameter set `model_1_ptm` was used for all experiments.

### C. Rosetta validation

For a subset of examples in all design studies, we validated designed proteins using the Rosetta suite of protein design and structure prediction tools [45]. We used fragment-assembly-based *ab initio* structure prediction [83] as an independent baseline for designed protein structures.

#### a. Ab initio structure prediction

Protein secondary structures were predicted using PSIPRED [84] and S4PRED [85] to provide secondary structures for fragment selection. 3-mer and 9-mer fragments were selected from the Rosetta fragment database based on secondary structure and sequence information. Alignment information was not used as designed sequences did not have sufficient homology to natural sequences. *Ab initio* structure prediction was performed by fragment-assembly using the AbinitioRelax protocol. Two starting conformations were evaluated: starting from an extended conformation and starting from the AF predicted structure. For both cases, 32000 decoys were generated. Decoys from both runs were pooled and scored using the Rosetta all-atom energy function [86]. The overall 10 lowest energy decoys were selected for relaxation. Decoys were relaxed using the Relax protocol. The AF predicted structure was also relaxed to generate 10 additional decoys in case *ab initio* failed to find a minimum energy structure. All decoys were rescored to compute directly comparable energies. The lowest energy relaxed decoy was selected as the final predicted structure and its Rosetta score and RMSD with respect to the AF predicted structure were evaluated.

#### b. Protein-protein interface analysis

As Rosetta does not allow for *ab initio* structure prediction of protein complexes other than in the case of symmetric homo-oligomers [87], interfaces of designed complexes were assessed using the InterfaceAnalyzer protocol [88, 89]. Structures were relaxed and Rosetta binding energy was computed by removing a single monomer from the complex, followed by repacking. As a further measure of interface quality, packing statistics were computed using the PackStat protocol [90], which detects solvent-inaccessible unfilled regions within an interface.

### D. Molecular dynamics simulation-based validation

A subset of designed monomers and higher-order complexes were further rigorously validated by performing all-atom molecular dynamics (MD) simulations in explicit solvent and analysing the corresponding properties of structural flexibility and internal/intramonomeric and interfacial contacts (for multimeric proteins and protein complexes).

#### a. Initial system construction

The standard AMBER force field (ff14sb) was used to describe all protein parameters [91]. Each protein or protein complex was solvated using atomistic TIP3P water [92] with a minimum of 10 Å of padding to form a cubic periodic box and then electrically neutralized with an ionic concentration of 0.15 M NaCl that used standard ionic parameters [93].

#### b. Simulation protocol

A standardised minimisation, equilibration and simulation protocol consisting of 11 stages was developed for all systems. A set of restraints (RS) were applied to each system at specified stages of equilibration. These consisted of restraining all heavy (non-hydrogen) atoms of the proteins. Each system was subsequently minimised across four stages with 1500 steps (500 steepest-descent + 1000 conjugate-gradient) of minimization applying restraints RS with different force constants in each sequential stage: Stage 1: 10 kcal/molA^2^, Stage 2: 5 kcal/molA^2^, Stage 3: 1kcal/molA^2^, Stage 4: unrestrained. MD simulations were performed in all subsequent stages. The SHAKE algorithm was employed on all atoms covalently bonded to a hydrogen atom. A time-step of 2fs was used. The long-range Coulomb interaction was handled using a GPU implementation of the particle mesh Ewald summation method (PME) [94, 95]. A nonbonded cutoff distance of 10 Å was used. In Stage 5, each system was heated from 10K to 300K in 1 ns and with RS (*k* = 10 kcal/molA^2^). The temperature was subsequently maintained at 300K using a Langevin thermostat with a damping constant of *γ* = 5.0 ps^-1^ and in Stage 6 the systems equilibrated for 1ns at constant volume, thus in the NVT ensemble. Subsequently the pressure was maintained at 1 atm using a Berendsen barostat with a pressure relaxation time of *τ_p_* = 1.0 ps and the systems simulated in the NPT ensemble for 100 ps for each of the subsequent stages with RS: Stage 7: *k* = 10 kcal/molA^2^, Stage 8: *k* = 5 kcal/molA^2^, Stage 9: *k* = 1 kcal/molA^2^, Stage 10: *k* = 0.5 kcal/molA^2^. Finally in Stage 11, RS restraints were removed and the systems simulated in the NPT ensemble for a further 5 ns. Following this, a production simulation of 100 ns each was performed for each system in the NPT ensemble with the same conditions as Stage 11. Coordinate snapshots from production simulations were generated every 10 ps, resulting in a trajectory of 10,000 snapshots per system.

#### c. Structural flexibility and contact analysis

Structural stability and flexibility were analysed by computing the root-mean squared deviation (RMSD) of the backbone atoms of the protein with respect to the initial predicted structure (after aligning the protein backbone) as well as the root-mean-squared fluctuation (RMSF) with respect to the average structure in the production MD. In the case of complexes, this was carried out both on the overall complex and for individual monomers separately. The number of sidechain-sidechain intramonomer contacts were computed for each snapshot of the production MD, based on a heavy atom distance threshold of 4Å. Similarly, interfacial contacts were computed for each interface in simulated protein complexes based on the same criteria. An overall picture of protein contacts was provided by summing the total intramonomer and interfacial contacts. Finally, for all simulated systems, an approximate potential of mean force (G) was determined by computing the Boltzmann-weighted distribution (*G* = –*k_B_T*ln(*ρ*)) in the 2D collective variable space of the global RMSF and total contacts (total intramonomer + total interfacial) contacts from the last 50 ns of simulation, where *ρ* is the normalised frequency in the binned 2D landscape.

### E. Structural clustering

Designed proteins of length 64 and larger were subdivided into a dictionary of fragments using Geometricus [96]. Fragments were collected using the *k*-mer method with *k* = 16 and the radius method with cutoff 10Å. A bag of features representation was computed for all designed proteins and dimensionality was reduced using non-negative matrix factorisation (NMF) with 50 components [97]. Proteins were separated into 10 clusters using ward-linkage agglomerative clustering [98] on their NMF components.

## IV. RESULTS

### A. AlphaFold predicts the structure of protein complexes

As recently shown by [18, 19], AF as trained on single-chain protein structures can be used for protein complex structure prediction. To gain confidence in the suitability of AF to design protein complexes using AlphaDesign, we first predicted the structure of small protein dimers, trimers, tetramers and pentamers (Fig. 2, SI Tab. S1). For the complexes considered, AF predictions show low root-mean-squared deviation (RMSD) to the native structure < 3 Å. While this may be due to these structures being part of the PDB [3] and thus part of AF’s training dataset [16], together with recent work on protein complex prediction with AF [18, 19], it provides some indication that AF could be suitable for the design of protein complexes using our framework.

**FIG. 2.**
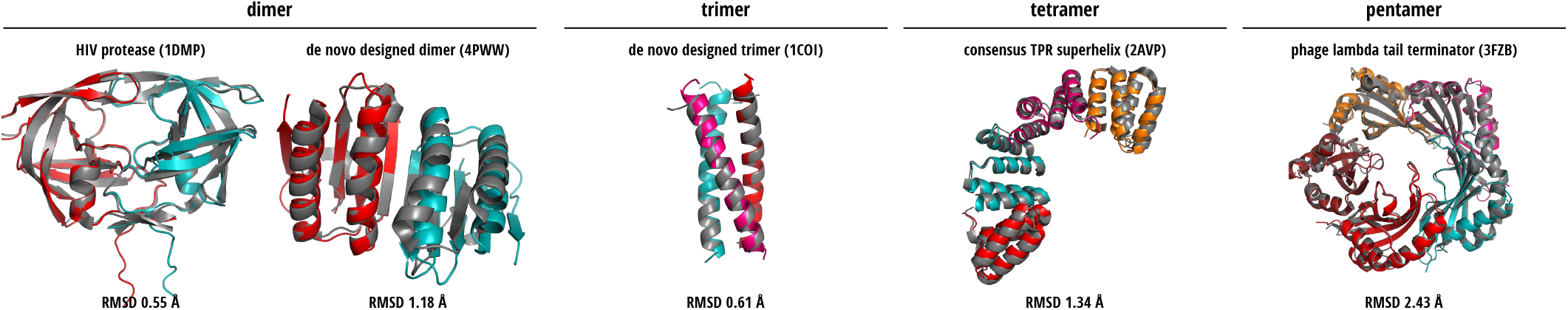
AlphaFold prediction of protein complexes. AlphaFold predictions of dimers (HIV protease 1DMP [99], de novo designed protein 4PWW [100]), trimers (de novo designed coiled coil 1COI [101]), tetramers (consensus TPR superhelix 2AVP), pentamers (phage lambda tail terminator 3FZB [102]). Predictions (colored) are aligned and overlaid onto the native structure (grey). All predictions show low RMSD to the the native structure.

### B. Single-sequence optimisation designs diverse monomers

As an initial step towards AF-based *de novo* protein design, we optimised protein sequences of length 32, 64, 128 and 256 amino acids to design monomeric, globular proteins (Fig. 3). All designed monomers show a high predicted aligned confidence (Fig. 3, predicted error), all exceeding a mean predicted aligned confidence of 0.82 and mean pLDDT of 0.83 (normalised to the range [0, 1]). This indicates that AF assigns high confidence to the predicted structure, as pLDDT of 0.7 or higher corresponds to a confident prediction of backbone structure [16, 17]. To validate the AF-predicted structure, we performed fragment-based *ab initio* structure prediction using Rosetta [45] for a subset of designs, starting from an extended amino acid chain, as well as from the AF structure (SI Fig. S1). For each monomer, we then extracted the 10 lowest-energy Rosetta structures. We relaxed these structures as well as the AF structure using the Rosetta force field [86] and chose the lowest-energy structure. For most monomers, the RMSD between the AF structure (grey, Fig. 3 (A-D), structures) and the lowest-energy structure (red) is less than 3.0 Å. For structures with distances exceeding this threshold (Fig. 3, C: result 8) the larger deviation seems to be due to a flexible *α*-helix and the AF predicted structure has a comparably low Rosetta energy.

**FIG. 3.**
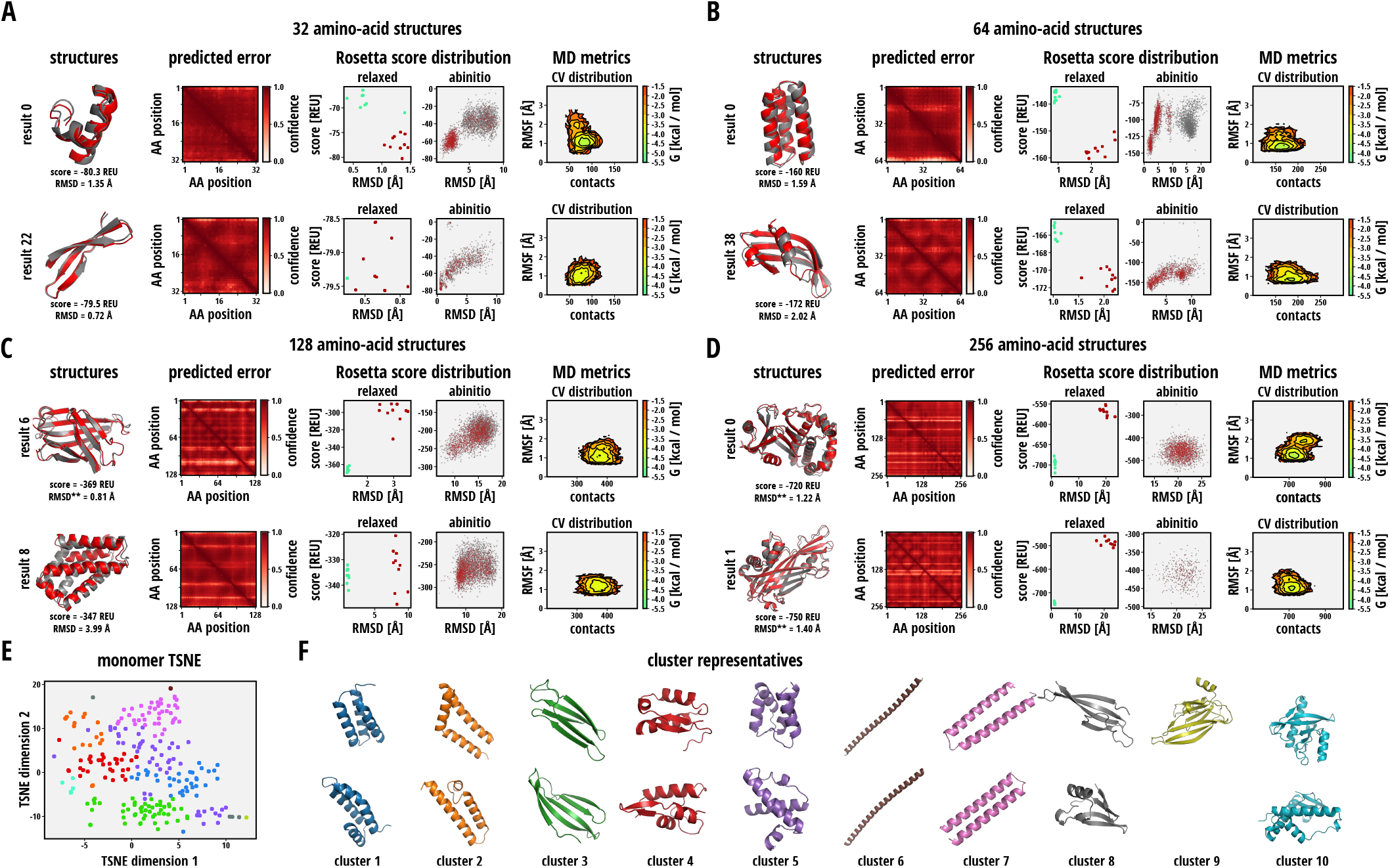
De novo monomer design. **(A - D)** *ab initio* and molecular dynamics validation of designed monomers of length 32 - 256 amino acids. Designed monomers (grey) are overlaid with their lowest-energy structure (red) from Rosetta *ab initio* prediction or relaxation of their predicted all-atom structure (structures). Each monomer is reported with its RMSD with respect to the AlphaFold structure and Rosetta energy. Predicted aligned confidence (pAC) is shown for each designed monomer (predicted error). For validation using Rosetta, the distribution of Rosetta scores and RMSDs to the AlphaFold structure is shown (Rosetta score distribution): for the top 10 relaxed structures (relaxed) starting from *ab initio* prediction (red) and the AlphaFold structure (blue); for all decoys from Rosetta *ab initio* prediction (ab *initio)* starting from an extended conformation (grey) and the AlphaFold structure (red). For molecular dynamics validation (MD metrics), the Boltzmann-weighted collective variable (CV) distribution in the 2D landscape of the number of intramonomer amino acid contacts and backbone RMSF of the protein with respect to the averaged MD structure is shown (CV distribution). **(E)** TSNE of designed monomer structures of length 64 amino acids and larger. Structures are separated into 10 clusters using agglomerative clustering. **(F)** Representatives for each cluster, showing diverse structures designed using AlphaFold.

To visualise the energy landscape of folds for each designed monomer, we inspected the distribution of decoys from *ab initio* folding with their Rosetta energy [86] and RMSD to the AF structure (Fig. 3 (A-D), Rosetta score distribution). We note that for monomers of size 32 and 64 amino acids, relaxations of the predicted structure (blue) have comparable, yet higher energies compared to the best 10 decoys from *ab initio* prediction (red) (Fig. 3 (A, B), Rosetta score distribution, relaxed). Strikingly, for monomers of size 128 and 256, relaxed AF structures (blue) exhibit far lower energies compared to *ab initio* decoys (red) (Fig. 3 (C, D)). This indicates a failure of *ab initio* structure prediction to find a reasonable minimum energy structure for those proteins. Indeed, the distribution of decoys for monomers of size 32 and 64 shows funnel shaped distributions with a single minimum at both low energy and low RMSD characteristic for protein folding (Fig. 3 (A, B), Rosetta score distribution, abinitio). In contrast, for sizes 128 and 256, the distribution is much more diffuse with no clear minimum being visible, indicating failure of *ab initio* prediction to find a reasonable minimum (Fig. 3 (A, B), Rosetta score distribution, abinitio). However, the observation that relaxed AF structures exhibit very low Rosetta energy [86], increases confidence that designed structures are plausible.

As a further rigorous validation step, we assess the structural stability of designed monomers by performing allatom molecular dynamics (MD) simulations. The Boltzmann-weighted frequency distribution in the 2D collective variable (CV) landscape consisting of backbone RMSF and intramonomer contacts shows that most systems exhibit a well-defined unimodal distribution centred on low RMSF and a significant number of contacts. This suggests the vast majority of conformers within the production ensemble sample a narrow range consistent with a maintenance of structural stability (Fig. 3 (A-D), MD metrics). Furthermore, the time evolution of the backbone RMSD with respect to the initial AF-predicted structure and backbone RMSF with respect to the MD-averaged structure are stably below 4Å and 2Å respectively in the majority of monomer systems (SI Fig. S1). Similarly, the time evolution of intramonomer contacts remains stable for all simulated monomers, suggesting a maintenance of folded structure. As expected the number of contacts grows with monomer size ranging from ~50-100, ~100-200, ~300-400 and ~600-800 for monomer lengths 32, 64, 128 and 256 amino-acids respectively (SI Fig. S1).

To assess the diversity of structures generated by AlphaDesign, we cluster all monomers of size 64 amino-acids or larger and extract representative structures for each cluster (Fig. 3 (E, F)). We embed structures using Geometricus [96], perform agglomerative clustering and visualise their distribution (Fig. 3 (E)). By visual inspection, cluster representatives encompass *α*-only (clusters 1, 2, 5, 6, 7), *β*-only (cluster 3) and mixed *αβ* proteins (clusters 4, 8, 9, 10) (Fig. 3 (F)).

### C. Searching for sequence pairs enables *de novo* dimer design

We next applied AlphaDesign to *de novo* design of protein homodimers and heterodimers. We optimised dimers with monomer size 32, 64 and 128 amino acids. As in our previous monomer designs, predicted dimers show high predicted aligned confidence for the AF structure (Fig. 4 (A-D), predicted error) with mean pAC > 0.75 and mean pLDDT > 0.83. By visual inspection, designed interfaces show a high level of shape complementarity. In lieu of *ab initio* structure prediction, which is in general harder to access for protein complexes using Rosetta [87], we opted for computing the Rosetta binding energy between monomers, as well as a score of interface packing statistics [90] as measures for structure quality. We applied this validation to a subset of designed dimers (SI Fig. S2). Dimers were relaxed using the Rosetta forcefield [86] noting that all relaxed structures have low RMSD with respect to the AF-predicted structure (Fig. 4, (A-F), complex structures). Furthermore, most dimers exhibit a packing statistic score greater than 0.6, which is comparable with the packing statistic of crystal structures with resolution 2.0 Å [90, 103]. Most structures show a Rosetta binding energy better than –40 REU hinting that predicted interfaces are stable under Rosetta [86].

**FIG. 4.**
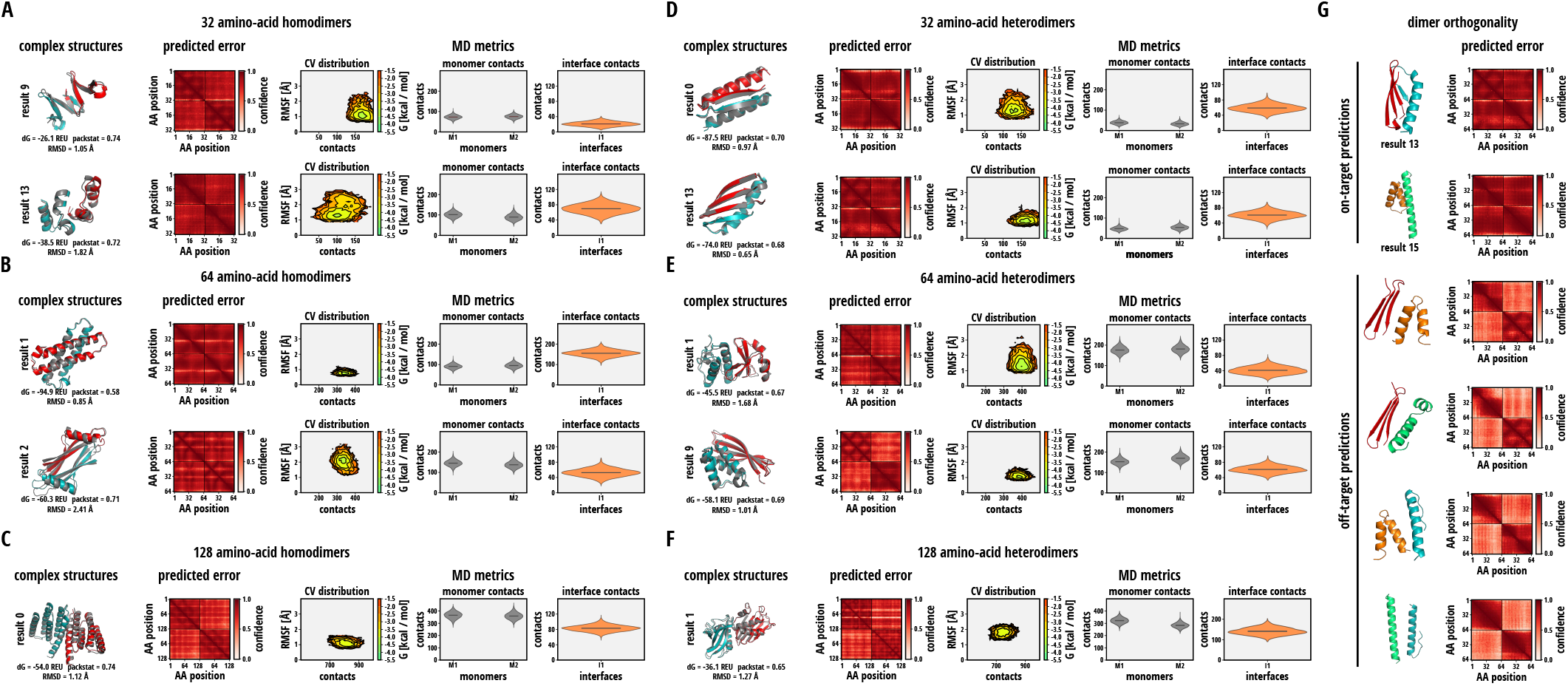
De novo dimer design. **(A - C)** Rosetta and molecular dynamics validation of designed homodimers of length 32 - 128 amino acids. Designed homodimers (grey) are overlaid with their lowest-energy structure (coloured) from Rosetta relaxation of their predicted all-atom structure (complex structures). Each relaxed dimer is reported with its RMSD to the AlphaFold structure, Rosetta binding energy and packing statistics. Predicted aligned confidence (pAC) is shown for each designed dimer (predicted error). For molecular dynamics validation (MD complex metrics), the Boltzmann-weighted CV distribution in the 2D space of total amino acid contacts (intramonomer+interfacial) and RMSF of the complex is shown (CV distribution), together with the distribution of individual monomer contacts and interface contacts. **(D-F)** Rosetta and molecular dynamics validation of designed heterodimers of length 32 - 128 amino acids. Measures displayed are the same as in (A-C). **(G)** Orthogonality of designed dimers. For two distinct pairs of designed dimers, predicted structures for the on-target and off-target complexes are shown. Predicted confidence (predicted error) of the on-target dimers is consistently higher for inter-monomer pairs of amino acids than the same confidence for off-target dimers.

MD simulations of dimer and higher order oligomer systems allow analysis of both intramonomer contacts as well as interfacial contacts between monomers. Again, the Boltzmann-weighted frequency distributions in the CV space of global RMSF and total contacts (intramonomer+interfacial) mostly show unimodal distributions with low mean RMSFs (≤ 2 Å) and significant numbers of total contacts that increase with complex size (Fig. 4, (A-F), MD metrics). Many dimer systems show stable time evolution of global RMSDs and RMSFs (SI Fig. S2), although significant deviation is observed in a few. Dissection of RMSDs, RMSFs, intramonomer and interfacial contacts, shows that both monomers in either the homodimer or heterodimer systems exhibit similar flexibility as well as number of intramonomer contacts. There are significant numbers of interfacial contacts in each dimer system, although, as expected these are fewer than the corresponding intramonomer contacts.

For synthetic biology applications, orthogonality is a desirable property for dimers [38]. That is, pairs of monomers designed to dimerise should only bind their designed partner, not monomers of other designed dimers. As a preliminary test for orthogonality of dimers designed using AlphaDesign, we predicted all combinations of monomers for a pair of designed dimers of monomer size 32 amino-acids (Fig. 4 (G)). On-target complexes were predicted by AF with high confidence (Fig. 4, (G), on-target predictions). In contrast, for off-target complexes – that is, complexes of monomers not designed for dimer formation – the mean predicted aligned confidence for inter-monomer amino acid pairs dropped below 0.5. This indicates that AF cannot confidently predict these off-target combinations as a dimer, providing preliminary evidence that dimers designed using AlphaDesign can indeed exhibit orthogonality.

### D. Multiple-sequence optimisation allows for homo-oligomer design

To demonstrate the feasibility of protein complex design beyond dimers, we proceeded to design homo-oligomers from trimers to hexamers. Monomer sizes of 32 and 64 amino acids were considered for trimers, tetramers, pentamers and 32 amino acids for hexamers. As before, we evaluated AF predicted aligned confidence, Rosetta binding energy, packing statistics and behaviour under 100 ns of molecular dynamics simulations for a subset of oligomers (SI Fig. S3). Designed structures show high predicted aligned confidence with most structures at pAC and pLDDT > 0.7 (Fig. 5, (A-G) predicted error), low RMSD to the Rosetta relaxed structure < 2.0Å, good binding energy < –40REU and packing statistics > 0.59 comparable to natural protein complexes [90, 103] (Fig. 5 (A-G) complex structures).

MD simulations of homo-oligomers exhibit similar properties to the dimers, with time evolution showing generally modest RMSDs (≤ 6 Å) and RMSFs (≤ 2 Å) and significant numbers of intramonomer and interfacial contacts (SI Fig. S3). Boltzmann-weighted distributions in the RMSF-contact space again show unimodal distributions with mean RMSFs remaining around 2Å and total contacts scaling with complex size. Dissection of individual intramonomer and interfacial contacts shows notable contacts in each monomer and interface for almost all systems (Fig. 5 (A-G) MD metrics).

Interestingly, inspection of designed complexes reveals a subset of extended oligomers with exposed interfaces on both sides (Fig. 5 (G-H)). The designed sequences could potentially oligomerise into larger assemblies. Correspondingly, MD simulations of these complexes show maintenance of intramonomer contacts for all monomers and sequential interfacial contacts for all but one interface, as expected. This indicates the possibility of designing and validating large-scale assemblies in AF beyond simple oligomers.

### E. Designing with a fixed monomer finds binders for target proteins

As a natural extension to dimer and oligomer design using AF with potential applications in biologics design and synthetic biology, we designed proteins binding to a fixed target protein (Fig. 6, SI Fig. S4). Here, we chose two proteins of length 64 amino-acids previously designed as target proteins and which exhibit distinctly different folds (Fig. 6 (A, B) target protein). We fixed the sequence for the target proteins during the design process while optimising the sequence of the designed binding proteins. Predicted structures of designed binding proteins exhibit high aligned confidence > 0.81, good Rosetta binding energy < –58REU and packing statistics > 0.67 comparable to natural protein interfaces [90, 103] (Fig. 6 (A, B)). This indicates that designed binders form dimers with the target proteins. By visual inspection we note that binders designed for each target protein seem to all bind at the same interface indicating preferences for binding sites during the design process.

**FIG. 5.**
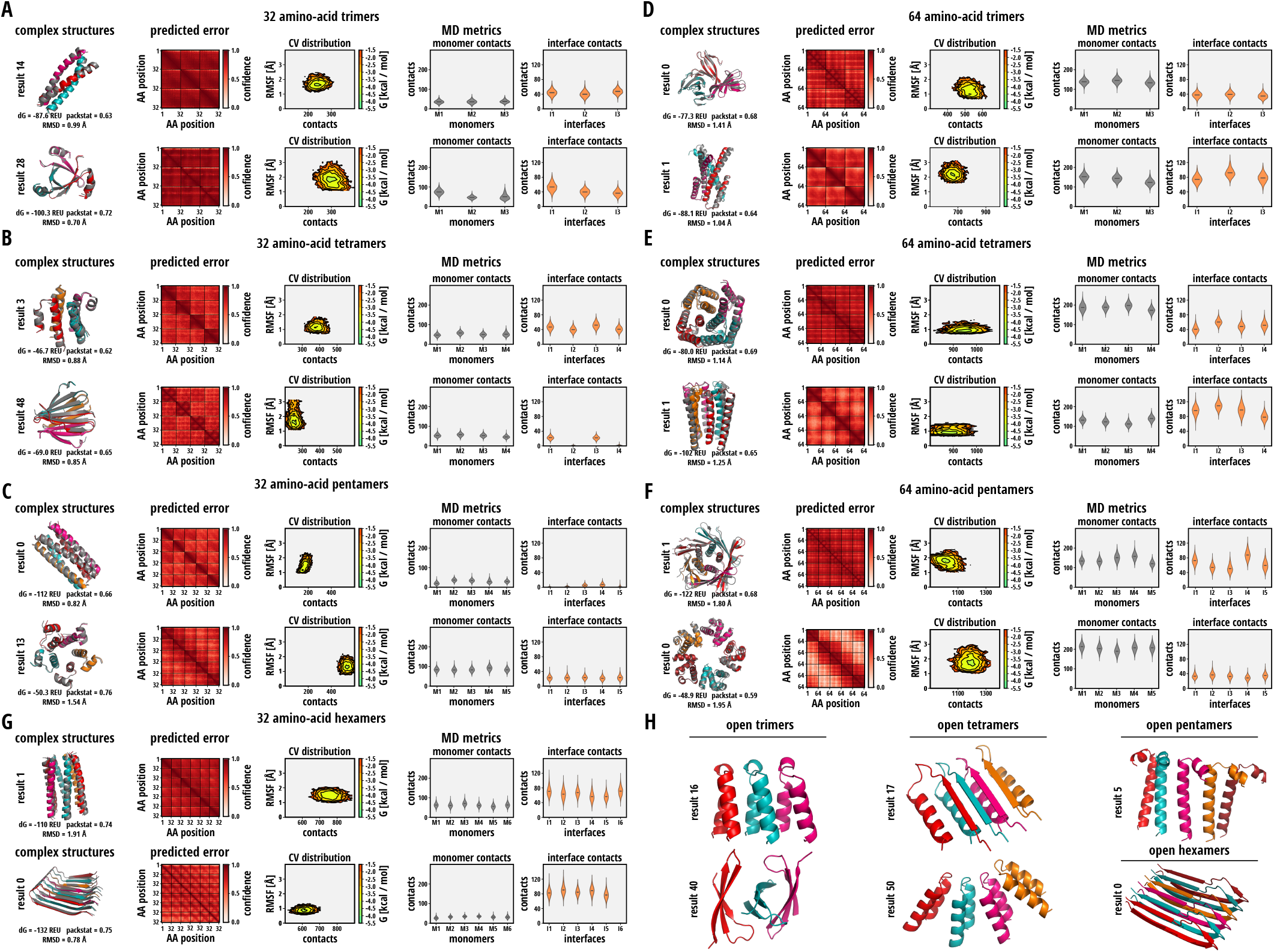
De novo oligomer design. **(A - G)** Rosetta and molecular dynamics validation of designed trimers, tetramers and pentamers of monomer length 32 - 64 amino acids, as well as hexamers of monomer length 32 amino acids. Designed oligomers (grey) are overlaid with their lowest-energy structure (coloured) from Rosetta relaxation of their predicted all-atom structure (complex structures). Each relaxed oligomer is reported with its RMSD with respect to the AlphaFold structure, Rosetta binding energy and packing statistics. Predicted aligned confidence (pAC) is shown for each designed oligomer (predicted error). For molecular dynamics validation, the Boltzmann-weighted CV distribution in the 2D space of total amino acid contacts (intramonomer+interfacial) and RMSF of the complex is shown (CV distribution), together with the distribution of individual monomer contacts and interface contacts. **(H)** Additional structures of extended oligomers. These oligomers can be extended by additional monomers on each side.

**FIG. 6.**
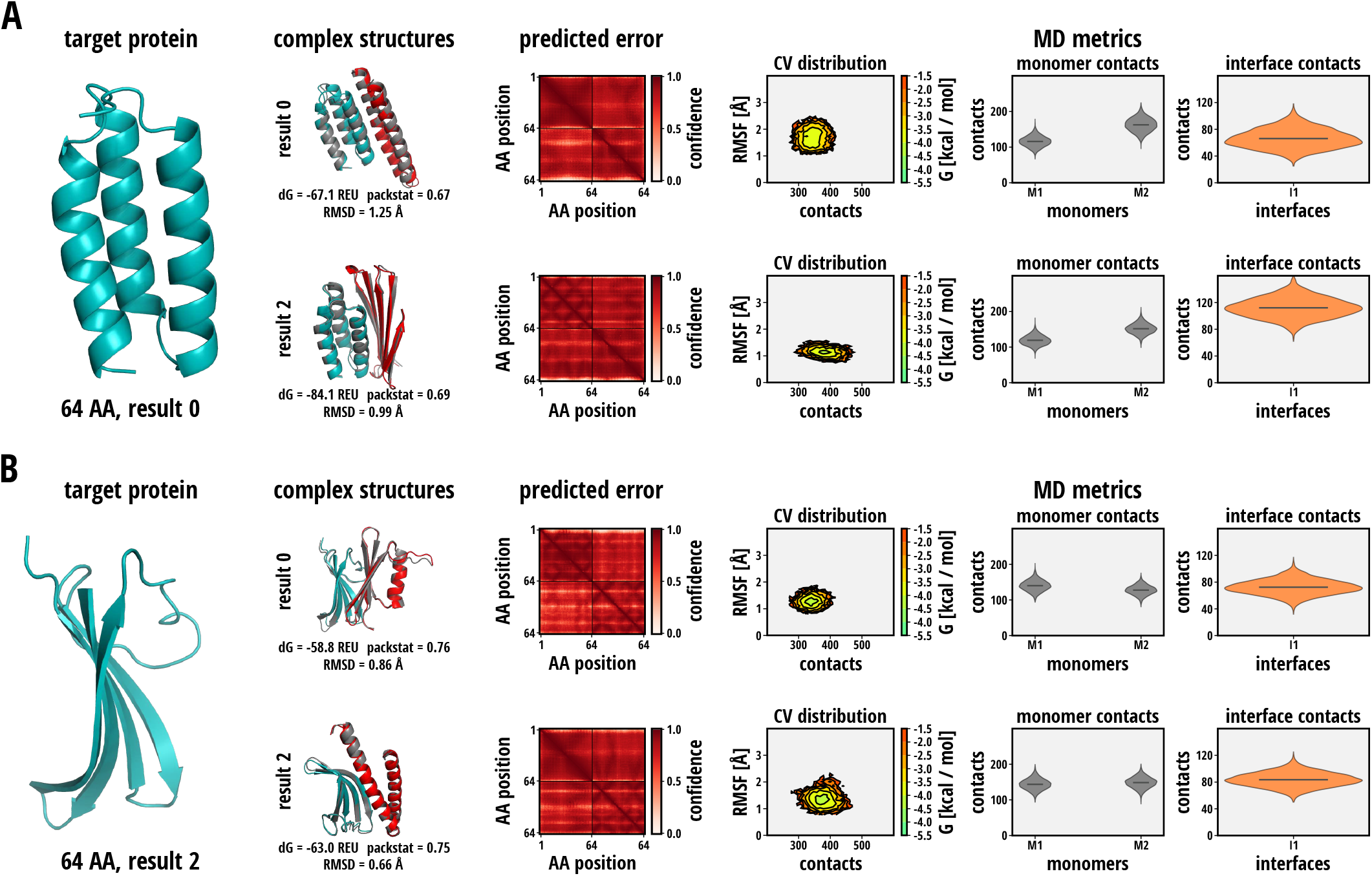
De novo binder design. **(A, B)** Rosetta validation of designed binders for previously *de novo* designed proteins. Relaxed binders (red) bound to the target protein (blue) are overlaid with their predicted all-atom structure (grey) (complex structures). Each relaxed binder is reported with its RMSD with respect to the AlphaFold structure, Rosetta binding energy and packing statistics. Predicted aligned confidence (pAC) is shown for each designed binder (predicted error). For molecular dynamics validation, the Boltzmann-weighted CV distribution in the 2D space of total amino acid contacts (intramonomer+interfacial) and RMSF of the complex is shown (CV distribution), together with the distribution of individual monomer contacts and interface contacts.

MD simulations of the two designed binders for each of the two target proteins show unimodal distributions in the 2D CV space of global RMSF-total contacts, with mean RMSFs < 2Å and mean number of total contacts ranging from 300-400, consisting of significant numbers of individual intramonomer (100-200) and interfacial (40-120) contacts.

### F. Sequence design uncovers signatures of conformational change in AlphaFold

To ascertain whether AF sequence space contains information about protein conformational change when interacting with other proteins, we predict the structures of two proteins known to have different monomeric and oligomeric states – KaiB, a circadian clock protein [104] and amyloid-*β* involved in Alzheimer’s disease [105]. As a monomer, KaiB adopts its ground-state structure, KaiBgs [104]. In complex with KaiC, KaiB changes conformation to the fold-switch stabilised state KaiBfs [104]. The AF prediction of KaiB as a monomer shows good agreement (RMSD 0.69 Å) with the native KaiBgs structure, while the prediction of KaiB in complex with KaiC shows good agreement with the native structure of KaiBfs (RMSD 1.92 Å). However, structural alignment of the predicted complex of KaiB and KaiC shows high RMSD (6.11 Å) to the structure of KaiBgs (Fig. 7 (A)), indicating that the AF prediction has captured a part of the conformational change between KaiBgs and KaiBfs. In its monomeric state, native amyloid-*β* forms an α-helical structure and changes conformation to a parallel *β*-sheet upon aggregation (Fig. 7 (A) grey) [105]. While not as accurate in terms of RMSD (> 5 Å), AF captures the transition between the *α*-helical monomeric state and the parallel *β*-sheet oligomer well (Fig. (A) red). This indicates that indeed AF may have learned to predict conformational change upon protein binding in a limited way.

**FIG. 7.**
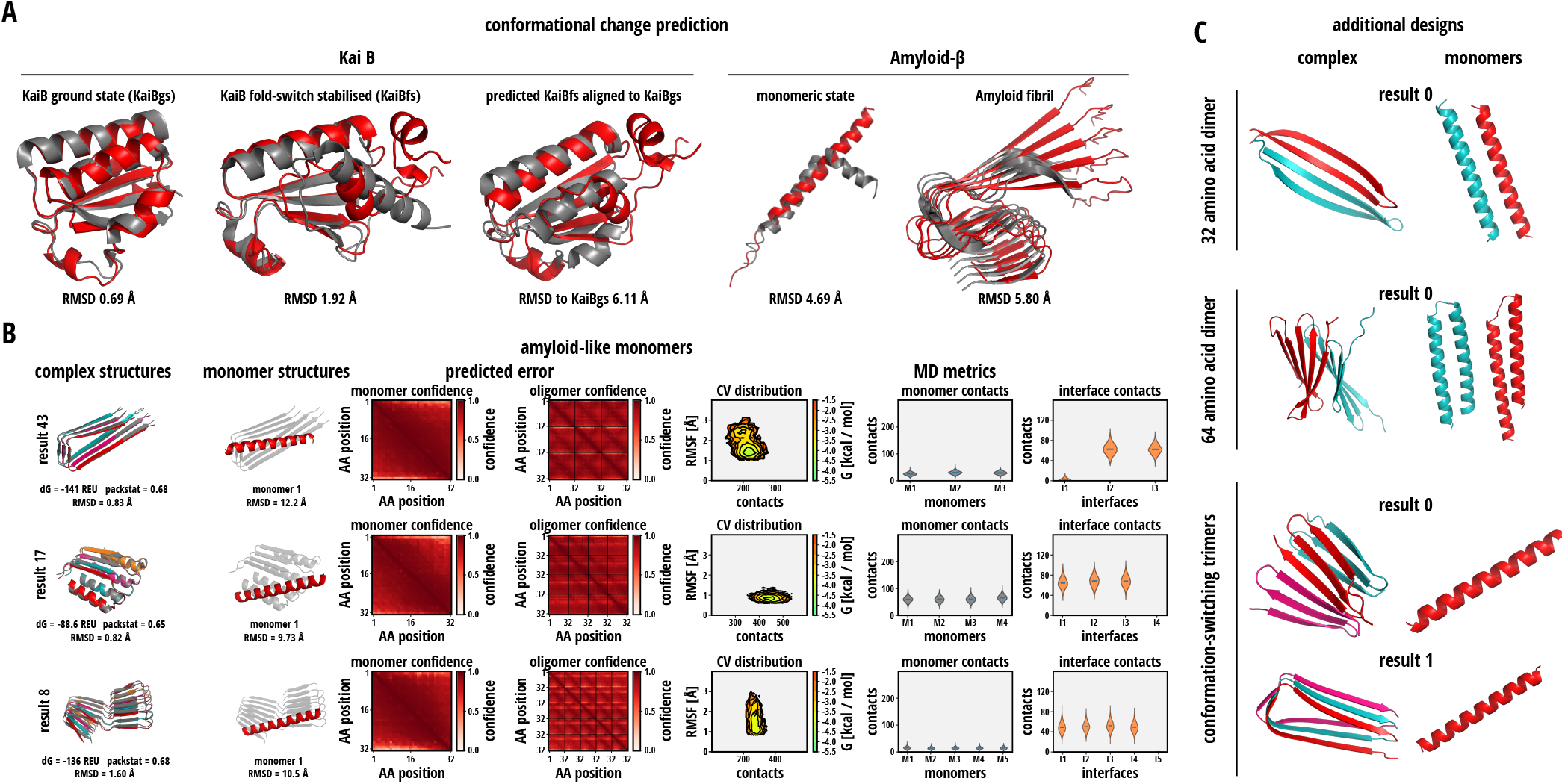
Conformational change in AlphaFold sequence space. **(A)** AlphaFold prediction of conformational change upon complex formation. (left) Predicted structures for monomeric circadian clock protein KaiB (KaiB ground state, KaiBgs) and fold-switch stabilised KaiB (KaiBfs) predicted in complex with KaiC (red) overlaid with the native structure of KaiBgs (1R5P [106]) and KaiBfs (5JYT [107]). Both predictions show low RMSD with respect to the corresponding native structure. Predicted KaiBfs (red) overlaid with native KaiBgs (the incorrect conformation, grey) shows high RMSD. (right) Predicted structures for monomeric Amyloid-*β* and its pentamer (red) overlaid with the native monomer (1IYT [108]) and oligomer (2MXU [109]). AlphaFold predicts the conformational change from *α*-helix to parallel *β*-sheet characteristic for amyloids. **(B)** Rosetta and molecular dynamics validation of designed oligomers of monomer length 32 showing amyloid-like predicted conformational change. Designed oligomers (grey) are overlaid with their lowest-energy structure (coloured) from Rosetta relaxation of their predicted all-atom structure (complex structures). Each relaxed oligomer is reported with its RMSD to the AlphaFold structure, Rosetta binding energy and packing statistics. Monomers are shown in comparison to the oligomer structure (monomer structures) to illustrate conformational change upon oligomerisation. Predicted aligned confidence (pAC) is shown for each designed oligomer and its constituent monomers (predicted error). For molecular dynamics validation, the Boltzmann-weighted CV distribution in the 2D space of total amino acid contacts (intramonomer+interfacial) and RMSF of the complex is shown (CV distribution), together with the distribution of individual monomer contacts and interface contacts. **(C)** Additional proteins designed using a target function favouring conformational change upon complex formation.

Furthermore, we identify a set of oligomer designs exhibiting conformational change upon complex formation (Fig. 7 (B), SI Fig. S5). Designed structures show a conformational change from an a-helix in the monomeric state (Fig. 7 (B), monomer structures) to a stack of parallel *β*-sheets characteristic for amyloids [105]. Both the monomeric and oligomeric state exhibit high predicted aligned confidence under AF (Fig. 7 (B) predicted error) and the oligomeric state shows binding energies < –88REU and packing statistics > 0.65 for all oligomers. This indicates that the complexes are stable under the Rosetta forcefield [86].

Similarly, MD simulations show well-defined minima in the RMSF-contact space centering on low RMSFs (< 2 Å) and a significant number of contacts (100-500). However, these conformation-switching open-ended oligomers do vary significantly compared to other oligomers including conformation-retaining open-ended oligomers, described previously. Whilst open-endedness is captured by a significant number of contacts in all-but-one sequential interfaces, the number of interfacial contacts (40-80) is either similar to or larger than the number of intramonomer contacts (10-60). This confirms the elongated conformational structure of monomers in the oligomeric state.

We next attempt to design heterodimers and homo-oligomers exhibiting conformational change upon complex formation. By maximising TM score between the monomeric and oligomeric state during optimisation, we find proteins exhibiting the desired conformation change (Fig. 7 (C)). Interestingly, we find the resulting proteins show conformational changes beyond the amyloid-like transition found in our previous design experiments, indicating a larger variety of conformation changing proteins present in AF sequence space.

## V. DISCUSSION AND CONCLUSIONS

Here, we have developed a *de novo* protein design framework based on sequence optimisation using evolutionary algorithms. Extending on previous works that utilise structure predictors[70, 71], we have embedded AlphaFold (AF) [16] into the design loop as a prediction oracle. Importantly, optimisation is facilitated by AF’s output predicted confidence measures, namely the predicted local distance difference test (pLDDT) [78] and the predicted aligned error (pAE) [16]. Furthermore, we have developed both a set of flexible target functions that encode various design tasks as well as an extendable platform for developing further target functions for solving bespoke design problems. This has enabled a range of applications including *de novo* design of protein monomers, dimers, oligomers, context dependent conformational switchers and binders to target proteins.

Our predicted structures are extensively validated using the Rosetta suite of protein design and structure prediction tools [45]. We also use fragment-assembly-based *ab initio* structure prediction [83] as an independent baseline for designed protein structures. In addition to this we develop a further rigorous validation protocol using all-atom molecular dynamics (MD) simulations that extends beyond conventionally used computational techniques for structure prediction evaluation. MD simulations enable extensive exploration of a putative native state, thus instabilities are picked up as increases in global structural flexibility, loss of internal contacts within protein monomers and/or loss of interfacial contacts within complexes.

We applied our framework to design *de novo* monomer proteins starting from completely random sequences ranging in size from 32 - 256 amino acids in length, based on a target function that combined pLDDT and pAE. This results in a range of structurally stable *de novo* designed monomer proteins with diverse folds. Using AF’s recently noticed functionality to predict complexes we incorporated and tested complex prediction on a number of systems showing good structural agreement. By specifying a further target function based on globular compactness together with complex prediction, we further designed a range of stable *de novo* protein complexes including homodimers, heterodimers and homo-oligomers from trimers to hexamers. Interestingly, we demonstrate orthogonality between pairs of dimers, suggesting the approach could have applicability in designing mutually exclusive combinations, for example, in protein logic gates [38]. Moreover, we have observed a number of open-ended predicted complexes, providing a potential route to the design of self-assembling systems [39].

A particularly intriguing subset of our oligomer systems exhibit striking conformational changes between their monomeric and oligomeric state. Structural prediction of existing protein systems known to change conformation and/or fold between monomer and oligomer form, including amyloid *α* – *β* switching, show that AF inherently contains signatures of conformational change. We consequently observe conformation switching between monomer and oligomer forms in some *de novo* designed open-ended oligomer systems. By developing a target function that maximises the structural difference between monomer and oligomer forms, our framework is able to *de novo* design oligomers with this conformational switching property. Context dependent conformational switching is a desirable feature in synthetic biology applications. For example, designing proteins to self-assemble inside but not outside the cell may be achievable by design of membrane permeable *α*-helices that spontaneously switch into membrane impermeable *β*-sheeted filaments as accumulated intracellular concentration drives an equilibrium towards the oligomeric state.

Finally, our approach enabled us to select a protein to be unmodified during the design loop - thus by combining this with a *de novo* designed protein starting from a random sequence we could design monomeric binders for a pre-specified target protein. Application of this to a set of target proteins resulted in the design of stable binders that exhibit significant interfacial contacts across the same interface for a given target protein. This suggests our framework could be further optimised towards therapeutic applications in potent biologic design.

## VI ACKNOWLEDGEMENTS

J.O.K. acknowledges support by the BMBF (de.NBI project: 031A537B) and the Volkswagen Foundation (contract 95826). S.K.S. acknowledges support by the Volkswagen Foundation “Experiment! Funding Initiative” (grant no. 93874-1).

## Notes

### Competing Interest Statement

The authors have declared no competing interest.

