## Supplementary Information for "AlphaDesign: A *de novo* protein design framework based on AlphaFold"

### AlphaDesign: A *de novo* protein design framework based on AlphaFold
### Supplementary Information

Michael Jendrusch ,* Jan O. Korbel ,† and S. Kashif Sadiq ‡

*Genome Biology Unit, European Molecular Biology Laboratory (EMBL), Meyerhofstrasse 1, Heidelberg 69117, Germany*

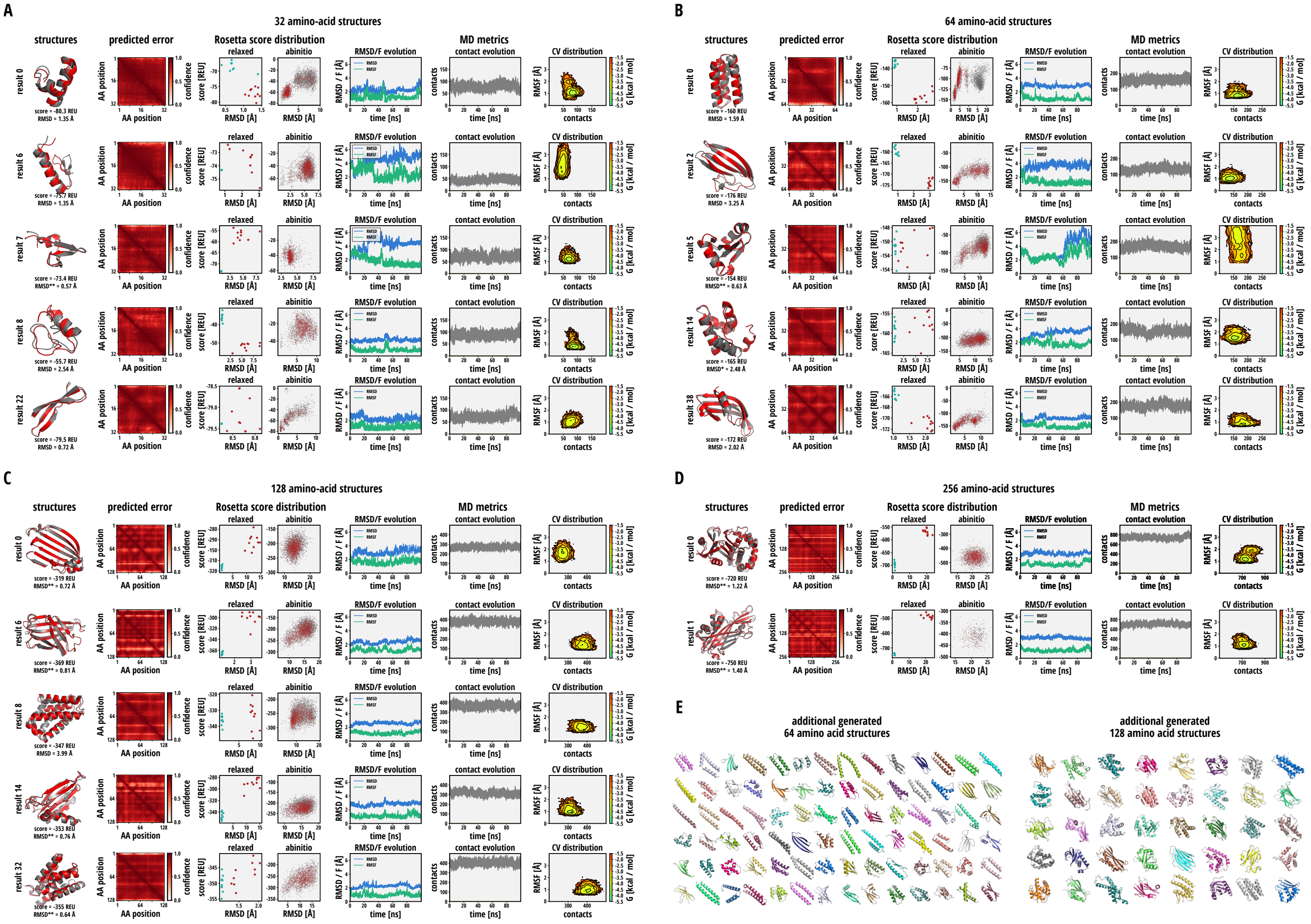


FIG. S1. Designed monomer validation using Rosetta and molecular dynamics. (A - D) Validation of *de novo* designed monomers of length 32, 64, 128 and 256 amino acids using Rosetta and molecular dynamics. (structures) The lowest Rosetta energy structure (red) is shown overlaid with the predicted AlphaFold structure (grey), reporting RMSD and Rosetta all-atom score. (predicted error) shows the predicted aligned confidence for each monomer. (Rosetta score distribution) shows the distribution of decoys from relaxation and *ab initio* prediction over Rosetta energy and RMSD to the predicted AlphaFold structure. (*relaxed*) shows this distribution for relaxations of the 10 lowest-energy *ab initio* decoys (red) and 10 relaxations of the AlphaFold predicted structure (blue). (*ab initio*) shows the distribution of decoys with Rosetta energy $< 0$ REU sampled starting from an extended conformation (grey) and the AlphaFold structure (red). For molecular dynamics-based validation, (RMSD/F evolution) shows the time evolution of RMSD (blue) and RMSF (green) for each monomer over the course of 100 ns of explicit-solvent molecular dynamics simulation. (contact evolution) shows the time evolution of the number of intra-monomer contacts over 100 ns of simulation (grey). (CV distribution) shows the Boltzmann-weighted distribution in the 2D collective variable (CV) landscape of backbone RMSF and total contacts (intramonomer+interfacial) for all snapshots in the last 50 ns of simulation. (E) additional random designed structures for monomers of length 64 and 128 amino acids.

---

*

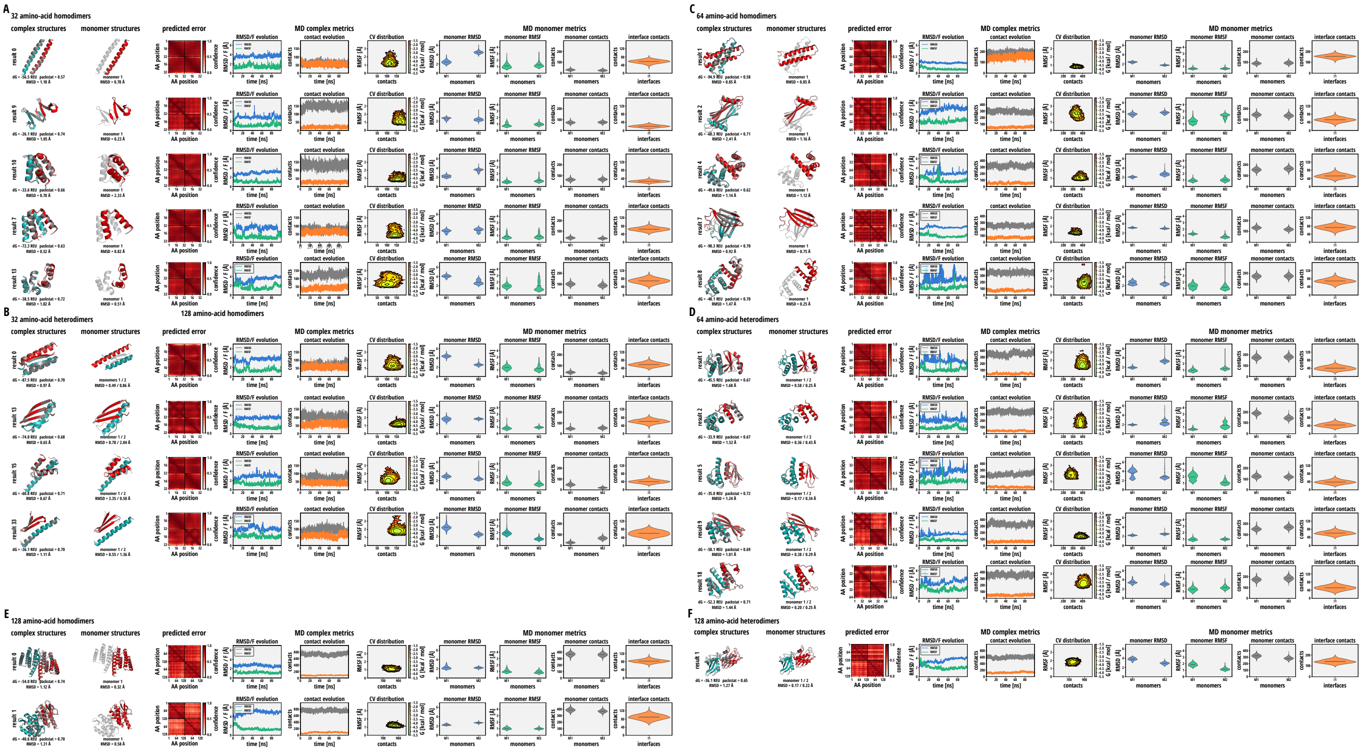

FIG. S2. Designed dimer validation using Rosetta and molecular dynamics. (A-F) Validation of *de novo* designed homo and heterodimers of monomer length 32, 64 and 128 amino acids using Rosetta and molecular dynamics. (complex structures) shows the AlphaFold predicted structure (grey) overlaid with the best of 10 relaxations (coloured) using the Rosetta all-atom score function. Structures are reported with their RMSD to the relaxed structure, as well as Rosetta binding energy $dG$ and packing statistics *packstat*. (monomer structures) shows the AlphaFold predicted structure of the monomer (red) overlaid with the structure of the complex (grey) and reports aligned RMSD. (predicted error) shows the predicted aligned confidence for each AlphaFold complex prediction. For molecular dynamics-based validation, (RMSD/F evolution) shows the time evolution of RMSD (blue) and RMSF (green) of the entire complex over the course of 100 ns of explicit-solvent molecular dynamics simulation. (contact evolution) shows the time evolution of total intra-monomer contacts (grey) as well as interface contacts (orange) over the course of 100 ns of simulation. (CV distribution) shows the Boltzmann-weighted distribution in the 2D collective variable (CV) landscape of backbone RMSF and total contacts (intramonomer+interfacial) for the entire complex for all snapshots in the last 50 ns of simulation. (MD monomer metrics) reports the distributions of RMSD (monomer RMSD, blue), RMSF (monomer RMSF, green), monomer contacts (grey) and interface contacts (orange) for all monomers and interfaces in the complex over the last 50 ns of simulation.

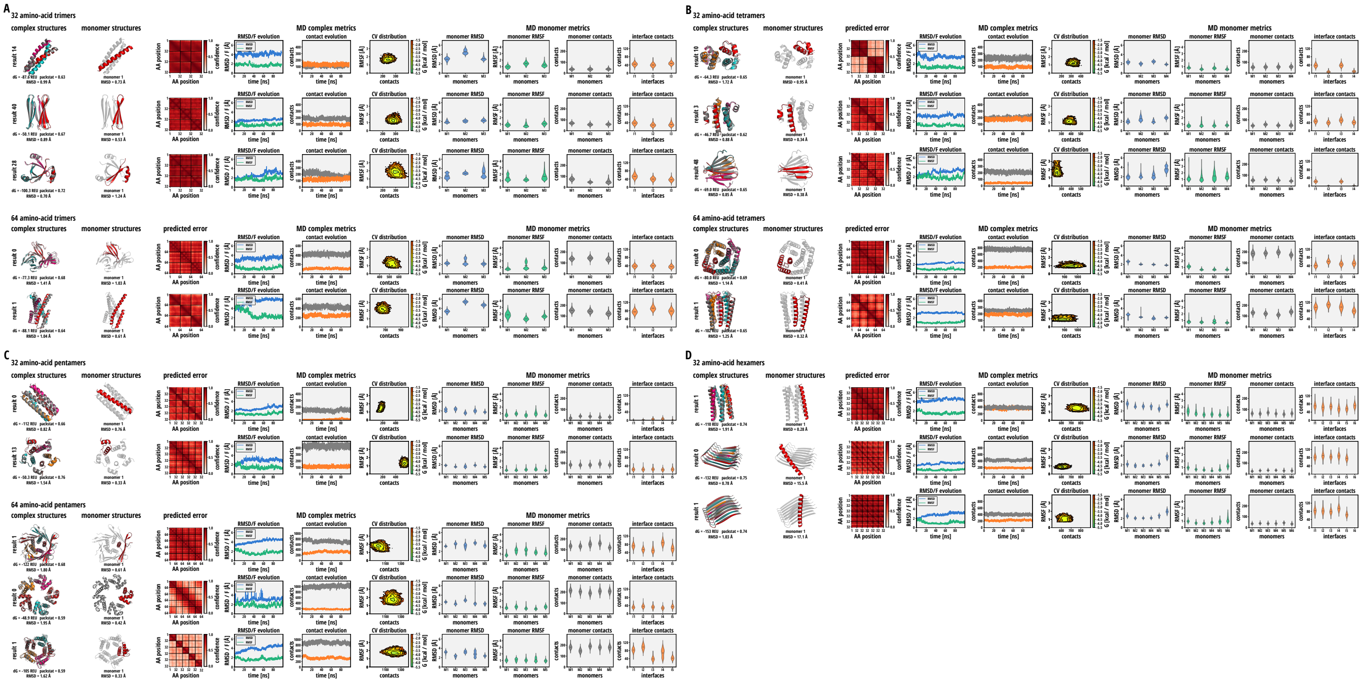

FIG. S3. Designed oligomer validation using Rosetta and molecular dynamics. (A-D) Validation of *de novo* designed trimers, tetramers, pentamers and hexamers of monomer length 32 and 64 amino acids using Rosetta and molecular dynamics. (complex structures) shows the AlphaFold predicted structure (grey) overlaid with the best of 10 relaxations (coloured) using the Rosetta all-atom score function. Structures are reported with their RMSD to the relaxed structure, as well as Rosetta binding energy of a single monomer to the rest of the complex $dG$ and packing statistics $packstat$. (monomer structures) shows the AlphaFold predicted structure of the monomer (red) overlaid with the structure of the complex (grey) and reports aligned RMSD. (predicted error) shows the predicted aligned confidence for each AlphaFold complex prediction. For molecular dynamics-based validation, (RMSD/F evolution) shows the time evolution of RMSD (blue) and RMSF (green) of the entire complex over the course of 100 ns of explicit-solvent molecular dynamics simulation. (contact evolution) shows the time evolution of total intra-monomer contacts (grey) as well as interface contacts (orange) over the course of 100 ns of simulation. (CV distribution)shows the Boltzmann-weighted distribution in the 2D collective variable (CV) landscape of backbone RMSF and total contacts (intramonomer+interfacial) for the entire complex for all snapshots in the last 50 ns of simulation. (MD monomer metrics) reports the distributions of RMSD (monomer RMSD, blue), RMSF (monomer RMSF, green), monomer contacts (grey) and interface contacts (orange) for all monomers and interfaces in the complex over the last 50 ns of simulation.

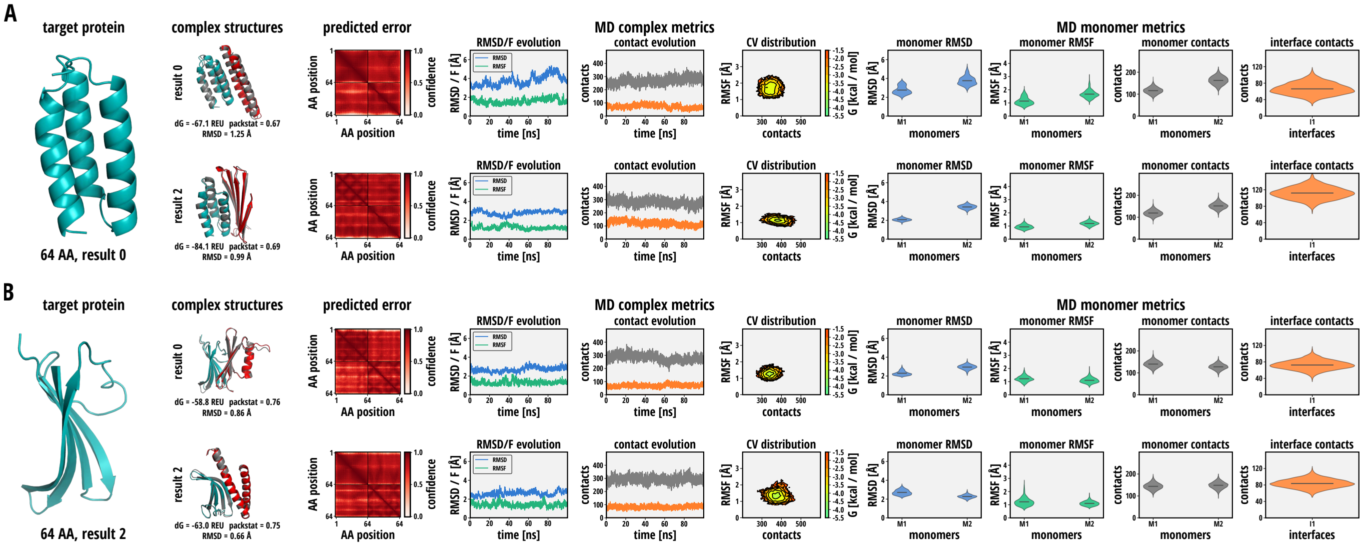


FIG. S4. Designed binding protein validation using Rosetta and molecular dynamics. (A, B) Validation of *de novo* designed binders of length 64 amino acids for two previously designed target proteins using Rosetta and molecular dynamics. (*target protein*) shows the AlphaFold predicted structure for the target protein. (complex structures) shows the AlphaFold predicted structure of the target protein in complex with the binder (grey) overlaid with the best of 10 relaxations (coloured) using the Rosetta all-atom score function. The target protein is coloured blue, all designed binders are coloured red. Structures are reported with their RMSD to the relaxed structure, as well as Rosetta binding energy $dG$ and packing statistics $packstat$. (monomer structures) shows the AlphaFold predicted structure of the monomer (red) overlaid with the structure of the complex (grey) and reports aligned RMSD. (predicted error) shows the predicted aligned confidence for each AlphaFold complex prediction. For molecular dynamics-based validation, (RMSD/F evolution) shows the time evolution of RMSD (blue) and RMSF (green) of the entire complex over the course of 100 ns of explicit-solvent molecular dynamics simulation. (contact evolution) shows the time evolution of total intra-monomer contacts (grey) as well as interface contacts (orange) over the course of 100 ns of simulation. (CV distribution)shows the Boltzmann-weighted distribution in the 2D collective variable (CV) landscape of backbone RMSF and total contacts (intramonomer+interfacial) for the entire complex for all snapshots in the last 50 ns of simulation. (MD monomer metrics) reports the distributions of RMSD (monomer RMSD, blue), RMSF (monomer RMSF, green), monomer contacts (grey) and interface contacts (orange) for all monomers and interfaces in the complex over the last 50 ns of simulation.

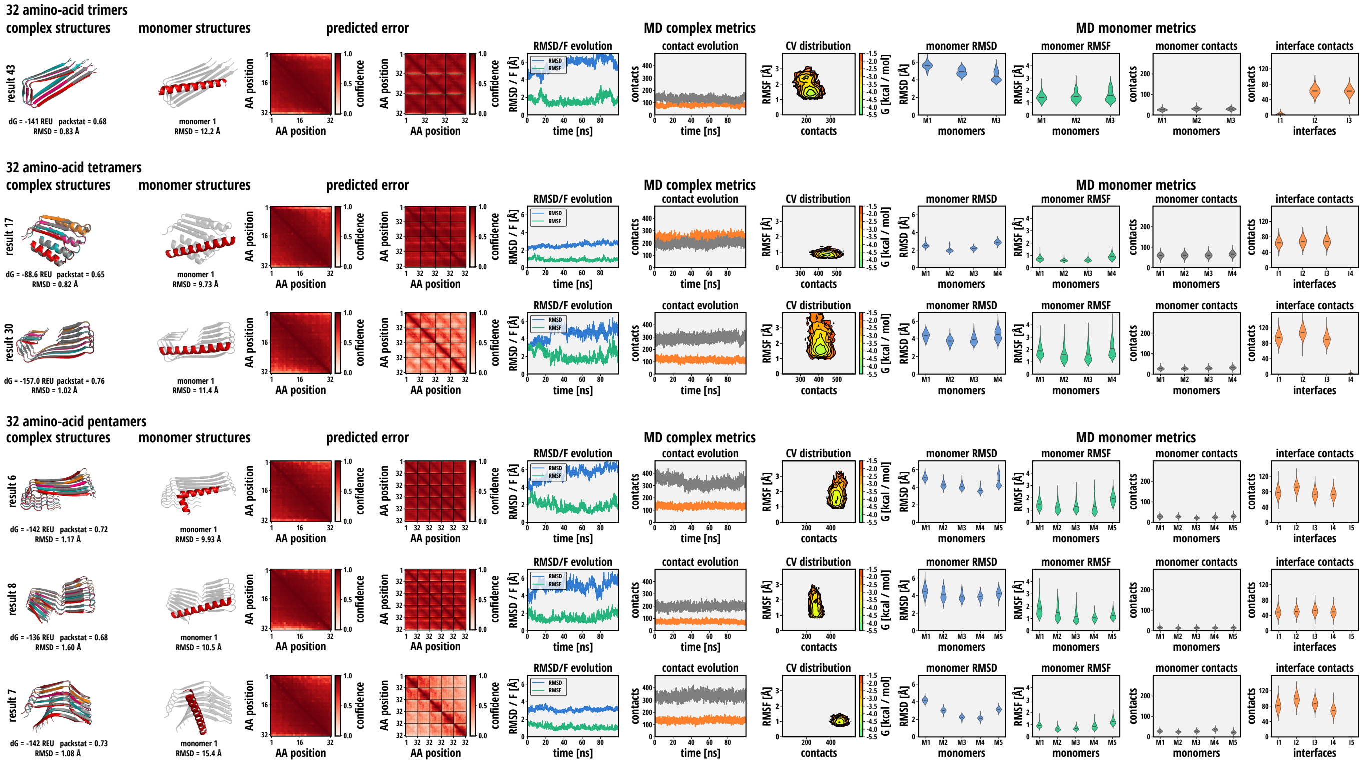


FIG. S5. Designed conformation-changing oligomer validation using Rosetta and molecular dynamics. (A-D) Validation of *de novo* designed trimers, tetramers and pentamers of monomer length 32 amino acids using Rosetta and molecular dynamics. (complex structures) shows the AlphaFold predicted structure (grey) overlaid with the best of 10 relaxations (coloured) using the Rosetta all-atom score function. Structures are reported with their RMSD to the relaxed structure, as well as Rosetta binding energy of a single monomer to the rest of the complex $dG$ and packing statistics $packstat$. (monomer structures) shows the AlphaFold predicted structure of the monomer (red) overlaid with the structure of the complex (grey) and reports aligned RMSD. (predicted error) shows the predicted aligned confidence for each monomer (left) and complex (right) AlphaFold prediction. For molecular dynamics-based validation, (RMSD/F evolution) shows the time evolution of RMSD (blue) and RMSF (green) of each complex over the course of 100 ns of explicit-solvent molecular dynamics simulation. (contact evolution) shows the time evolution of total intra-monomer contacts (grey) as well as interface contacts (orange) over the course of 100 ns of simulation. (CV distribution)shows the Boltzmann-weighted distribution in the 2D collective variable (CV) landscape of backbone RMSF and total contacts (intramonomer+interfacial) for the entire complex for all snapshots in the last 50 ns of simulation. (MD monomer metrics) reports the distributions of RMSD (monomer RMSD, blue), RMSF (monomer RMSF, green), monomer contacts (grey) and interface contacts (orange) for all monomers and interfaces in the complex over the last 50 ns of simulation.

| PDB ID | description | sequence input | homooligomer input | MSA method | $N$ recycling |
|---|---|---|---|---|---|
| 1DMP | HIV protease | VSFNFPQITLWKRPLVTIRIGGQLKEALLNTGADDTVLEEMNL<br>PGKWKPKMIGGIGGFIKVRQYDQIPVEICGHKAIGTVLVGPTP<br>VNIIGRNLLTQIGCTLNF | 2 | mmseqs2 | 48 |
| 4PWW | *de novo* designed homodimer | MEMDIRFRGDDLEALLKAAIEMIKQALKFGATITLSLDGNDLE<br>IRITGVPEQVRKELAKEAERLAKEFGITVTRTIRGSWSLEHHH<br>HHH | 2 | mmseqs2 | 48 |
| 1COI | *de novo* designed trimeric coiled coil | GEVEALEKKVAALESKVQALEKKVEALEHGG | 3 | single sequence | 48 |
| 2AVP | consensus TPR superhelix | GSAEAWYNLGNAYYKQGDYDEAIEYYQKALELDPRSAEAWYNL<br>GNAYYKQGDYDEAIEYYQKALELDPRS | 4 | mmseqs2 | 48 |
| 3FZB | phage lambda tail terminator | GSHMKHTELRAAVLDALEKHDTGATFFDGRPAVFDEADFPAVA<br>VYLTGAEYTGEELDSDTWQAELHIEVFLPAQVPDSELDAWMES<br>RIYPVMSDIPALSDLITSMVASGYDYRRDDDAGLWSSADLTYV<br>ITYEM | 5 | mmseqs2 | 48 |
| 1R5P | KaiB ground state (KaiBgs) | MAPLRKTYVLKLYVAGNTPNSVRALKTLNNILEKEFKGVYALK<br>VIDVLKNPQLAEEDKILATPTLAKVLPPPVRRIIGDLSNREKV<br>LIGLDLLYEEIGDQAEDDLGLE | 1 | mmseqs2 | 48 |
| 5JYT | KaiB foldswitch-stabilised (KaiBfs) in complex with KaiC | MAPLRKTYVLKLYVAGNTPNSVRALKTLNNILEKEFKGVYALK<br>VIDVLKNPQLAEEDKILATPTLAKVLPPPVRRIIGDLSNREKV<br>LIGLDLLYEEIGDQAEDDLGLE:<br>DYKDDDDKAEVKKIPTMIEGFDDISHGGLPQGATTLVSGTSGT<br>GKTLFAVQFLYNGITIFNEPGIFVTFEESPQDIIKNALSFGWN<br>LQSLIDQGKLFILDASPDPDGQEVAGDFDLSALIERIQYAIRK<br>YKATRVSIDSVTAVFQQYDAASVVRREIFRLAFRLAQLGVTTI<br>MTTERVDEYGPVARFGVEEFVSDNVVILRNVLEGERRRRTVEI<br>LKLRGTTHMKGEYPFTINNGINIFDYKDDDDK | 1:1 | mmseqs2 | 48 |
| 1IYT | peptide from amyloid-$\beta$, monomeric | DAEFRHDSGYEVHHQKLVFFAEDVGSNKGAIIGLMVGGVVIA | 1 | single sequence | 12 |
| 2MXU | peptide from amyloid-$\beta$, fibril | DAEFRHDSGYEVHHQKLVFFAEDVGSNKGAIIGLMVGGVVIA | 4 | single sequence | 12 |

TABLE S1. Parameters of structure-prediction runs using Colabfold.
